# Reduced expression of PER2 protein contrbutes to β_1_-AA induced cardiac autophagy rhythm disorder

**DOI:** 10.1101/2024.12.06.627296

**Authors:** Peng-Jia Li, Jia-Yan Feng, Jiao Guo, Jin Xue, Yang Li, Shi-Yuan Wen, Xiao-Hui Wang, Hui-Rong Liu, Li Wang

## Abstract

It has been confirmed that heart failure may be linked to fluctuations in autophagy rhythm of cardiomyocytes throughout the day. It is known that circadian rhythms depend on the regulation of core biological clock proteins, with PER2 playing a crucial role. Our previous research has confirmed the presence of β_1_-Adrenergic receptor autoantibodies (β_1_-AA) could induce inhibition of myocardial autophagy, leading to cell death and heart failure. However, it remains unclear whether β_1_-AA induces cardiac autophagy rhythm disorder by affecting PER2 expression. This study find that β_1_-AA disrupts the autophagy rhythm in cardiomyocytes, primarily indicates by the decreased expression of the autophagy marker protein LC3; β_1_-AA induces disruption of the rhythmic expression of PER2 protein in myocardial cells, mainly manifests by a decrease in PER2 protein expression; Metoprolol is employed to verify that the β_1_-adrenergic receptor contributes to the reduction of Per2 protein caused by β_1_-AA. Knocking down Per2 with lentivirus reduces the inhibition of LC3 expression caused by β_1_-AA, while overexpressing Per2 in cardiomyocytes using lentivirus significantly restores β_1_-AA-induced decline in LC3 expression. At the same time, mTORC1 activation is found to participate in β_1_-AA-induced autophagy inhibition of cardiomyocytes after pretreatment with the mTORC1 inhibitor rapamycin. Furthermore, it is confirmed that the decreased expression of PER2 protein caused by β_1_-AA disrupts the myocardial autophagy rhythm by promoting mTORC1 activation through lentiviruses that knock down or overexpress the Per2 gene. This study provides experimental basis for the precision treatment of cardiovascular diseases from the perspective of biological rhythm.

## Introduction

Heart failure is the final stage of various cardiovascular diseases ^[1]^.With the increasing aging of society, the prevalence and mortality of heart failure in the world are on the rise, of which China accounts for 1/3 ^[2, 3]^. Lifestyle related to circadian rhythm disturbance such as “daily staying up late”, “shift work” and “sleep disorders” can disrupt the repair cycle of the heart and cause damage or even death of cardiomyocytes, resulting in an increased risk of heart failure or aggravation of the pre-existing heart failure ^[4]^. Studies have shown that various biological activities that maintain the normal operation of cells have circadian changes, including autophagy ^[5]^. As we all know, cardia<u>c</u> adverse events occur frequently in the early morning, when autophagy activity is at a low level ^[6]^, suggesting that the occurrence of heart failure may be related to the circadian rhythm of autophagy in cardiomyocytes throughout the day. Therefore, it is particularly important to find the factors causing the disturbance of circadian autophagy rhythm in the process of heart failure.

Clinical data show that autoantibodies are present in the serum of 40%-60% of patients with heart failure, which continuously activate the β_1_-adrenergic receptor (β_1_-AR), namely the β_1_-adrenergic receptor autoantibodies (β_1_-AA) ^[7]^. Removal of β_1_-AA in serum of patients with heart failure by immunosorbent technique could significantly improve the cardiac function of patients ^[8]^. Our previous studies have shown that β_1_-AA can inhibit the level of myocardial autophagy and promote cardiac cell death leading to heart failure ^[9]^. However, it is not clear whether β_1_-AA is a factor causing the disturbance of cardiomyocyte autophagy rhythm.

Studies have found that autophagy rhythm is regulated by circadian rhythm ^[10]^. Circadian rhythm is a regulated rhythmic oscillation with a 24-hour cycle that exists universally in the body. Its molecular mechanism is a transcriptional translation feedback loop composed of a series of clock genes ^[11]^, in which the negative feedback regulation of PER2 protein plays a key role ^[12]^. Studies have found that Per2 is closely related to autophagy ^[13]^. After the silencing of Per2 gene in fibroblasts, the levels of autophagy related proteins LC3II and ULK1 decreased significantly, and their rhythms were also significantly abnormal ^[14]^. Therefore, we speculate that β_1_-AA could cause the disturbance of autophagy rhythm of cardiomyocytes by altering the expression of PER2 in cardiomyocytes, and further explore the specific mechanism from the perspective of mTORC1, in order to provide a new experimental basis for the precision treatment of cardiovascular diseases from the perspective of biological rhythm.

## Materials and Methods

### Animal models and Cells

C57BL/ Series 6 mice, male, 6 weeks to 8 weeks old, weighing 18 to 22 g/ mouse. SD rats, male, 6 weeks to 8 weeks old, weighing 180 g to 210 g/piece, all purchased from Laboratory Animal Center of Shanxi Medical University [Animal qualification Certificate No. : SCXK (Jin) 2015-0001]. H9c2 rat cardiomyocytes were purchased from Shanghai Cell Bank, Chinese Academy of Sciences.

### Experimental reagents and instruments

β_1_-AR-ECII antigen peptide, purchased from China Gill Biochemical (Shanghai) Go., LTD; DMEM medium (high sugar), purchased from Gibco, USA; The required one is purchased from abcam Company; The second resistance required was purchased from Zhongshan Jinqiao Company;dexamethasone from Solarbio; RNAiso plus, Prime Script RT Master Mix and TB Green probes were purchased from Baori Medical Technology Limited company. Cell incubator purchased from Eppendorf; Electrophoresis instrument, film transfer instrument and exposure instrument purchased from BIO-RAD company; Laser confocal from Olympus Corporation; Ultrasonic crusher purchased from the United States SONICS&MATERLALS company.

### Establishment of an active immune animal model

Male C57BL/6 mice aged 6 weeks to 8 weeks were randomly selected and divided into Control group and β_1_-AR active immune group according to random number table method.In the active immunization (β_1_-AA) group, β_1_-AR-ECII antigen mixed emulsion with a dose of 0.4 μg/g was injected into the subcutaneous back of mice, which was the first immunization, and the immunization was strengthened every 2 w for a total of 6 w. Solvent Control group: at the same time, the same dose of Na_2_CO_3_ and adjuvant mixed solution were injected into the subcutaneous back of mice, and the immunization time and frequency were the same as that of the active immunization group.

### SA-ELISA

The content of β_1_-AA in the serum of actively immunized mice was detected by SA-ELISA method, and the β_1_-AA in the serum of antibody positive mice was purified to prove that the active immunized mouse model of β_1_-AR was successful.The synthetic β_1_-AR-ECII antigen peptide was prepared into a solution and coated uniformly on the bottom of the microplate to produce a solid phase antigen.Isolation was carried out in sequence, incubation of serum to be tested, incubation of secondary antibody with biotin label, incubation of streptase lecitin with horseradase label and incubation of chromogenic solution. Finally, absorbance value was detected by enzyme marker.

### Concentration and purification of β_1_-AA

The rats were injected with the same active immunization method, and the serum of the successfully active immunized rats was mixed and concentrated using a special EP tube. Then the β_1_-AA in the serum was purified by affinity chromatography column, and its concentration was detected for subsequent cell experiments.

### Cell culture

The frozen H9c2 rat cardiomyocytes were resuscitated, the centrifuged cell precipitates were added to fresh medium for re-suspension, moved to a culture bottle and cultured in a 5% CO_2_ incubator at 37℃.When the cell condition is good, it can be treated with medicine.Control group: After dexamethasone treatment for 4 h, H9c2 cells were treated with purified negative IgG (1 μmol/l). β_1_-AA group: After dexamethasone treatment for 4 h, purified β_1_-AA was treated on H9c2 cells (1 μmol/l).

### Cell cycle flow cytometry detection

The cell cycle was measured by flow cytometry to reflect the degree of cell synchronization. H9c2 cardiomyocytes were uniformly spread in the six-well plate, and when the cell density reached 60% to 70%, dexamethasone (final concentration of 100 nM) was treated for 1 h, 2 h, and 4 h, respectively, and the cells could be collected and placed in a centrifugal tube containing 75% ethanol for low temperature fixation. Then, dye that can specifically bind to DNA was selected, and the cell suspension was absorbed and placed in the channel of flow cytometry for detection. Different fluorescence intensity showed in the results indicated different cell proportions at different stages. The proportion of the number of G0/G1 phase cells is an indicator to determine whether the cells have reached synchronization.

### Western blotting

The expression level of protein in myocardial tissue or cells was detected. First, the collected mouse tissue or cell samples were cleaved and the protein concentration was detected, and the samples were boiled according to the protein system (40 μg) in order to denature the protein. Then, polyacrylamide gel electrophoresis (SDS-PAGE) was carried out, and the cooked sample was added into the prepared glue hole, and the electrophoresis was stopped when the sample ran under the rubber plate.The PVDF membrane was activated with methanol, and the tape of appropriate size was cut to start the membrane transfer at 15 V for 15 min. PVDF membrane was immersed in milk solution and closed for 2 h, the first antibody was incubated at 4℃ overnight, and the second antibody was incubated for 2 h. The strips were coated with supersensitive luminescent solution and placed in the exposure instrument.Finally, Image software was used to analyze the exposed strips.

### Immunofluorescence

Immunofluorescence technique was used to detect the expression and localization of proteins in cells.After the cells successfully climbed the tablet, they were treated with drugs.It was fixed with paraformaldehyde, then the film was broken with TrizonX-100, sealed with 5% BSA, and the corresponding primary and fluorescent secondary antibodies were added (pay attention to avoid light).Finally, the protein fluorescence was observed by confocal laser after DAPI nucleation.

### CCK-8

100 μl cell suspension was inoculated into 96-well cell culture plates, and the number of cells per well was controlled at 20% ∼ 30%.The culture plate was placed at 37°C and incubated in 5% CO_2_ incubator for 24 h, then the PBS was cleaned and the liquid was changed, and a certain volume of the drug to be tested was added to the culture plate for 24 h.Finally, 10 μl CCK-8 reagent was added to each well, and after incubation in the incubator for 2 h, the absorbance value at 450 nm was measured using a multifunctional enzyme marker.The calculation formula is: relative cell viability (%) = (experimental group X value − blank control group X value)/(negative control group X value − blank control group X value) ×100%.The percentage obtained from each set of data is presented as cell survival.

### Statistical analysis

The SPSS 16.0 statistical package was used to calculate the mean ± standard deviation (x±s) of the data and generate a normal distribution curve.The difference of protein expression between the control group and the experimental group was analyzed by paired sample t test.At the same time, one-way analysis of variance was used to detect the independence of multiple sets of data.Values with a significance of *P*<0.05 were considered statistically significant.

JTK_CYCLE is a new nonparametric statistical algorithm for detecting and describing the rhythm portion of large-scale genomic data.JTK_CYCLE analysis was performed using the Windows -R 2.10.0 software package, 95% expected confidence interval was set, and JTK_*P* value in each group of output data was recorded (*P* value <0.05 after correction indicated that it had circadian fluctuations).

## Results

### 1. β_1_-AA disrupted myocardial autophagy rhythm, mainly by reducing the expression of autophagy marker protein LC3

In order to detect the endogenous autophagy rhythm of myocardial tissue, the mice were actively immunized with β_1_-AR-ECII, and the immunization was enhanced once every 14 days. After the first immunization, the mice in each group were placed in L/D environment (Light/Dark for 12 h) for 2 w. After the second immunization enhancement, the light source was turned off. Adjust to all black environment D/D (Dark/Dark 12 h) 2 w. CT0 is defined as the light on-set. SA-ELISA detection found that the OD value of β_1_-AA in serum of 4 w mice was significantly increased (Figure 1A), indicating that the model was successfully constructed. Western blot was used to detect LC3 protein expression in myocardium of mice in the Control group. The results showed that LC3II protein expression in myocardium of mice in the control group showed a significant circadian rhythm (JTK_CYCLE ADJ.*P*< 0.05) (Table 1). Compared with the Control group, LC3II protein expression in the myocardial tissue of actively immunized mice was significantly decreased at CT8 and CT12 levels, the total LC3 results were consistent with the LC3II trend (Figure 1B). The JTK_CYCLE analysis results showed that the rhythmic expression of LC3II protein was lost in the myocardial tissue of actively immunized mice (JTK_CYCLE ADJ.*P*>0.05) (Table 1).

**Fig 1.**
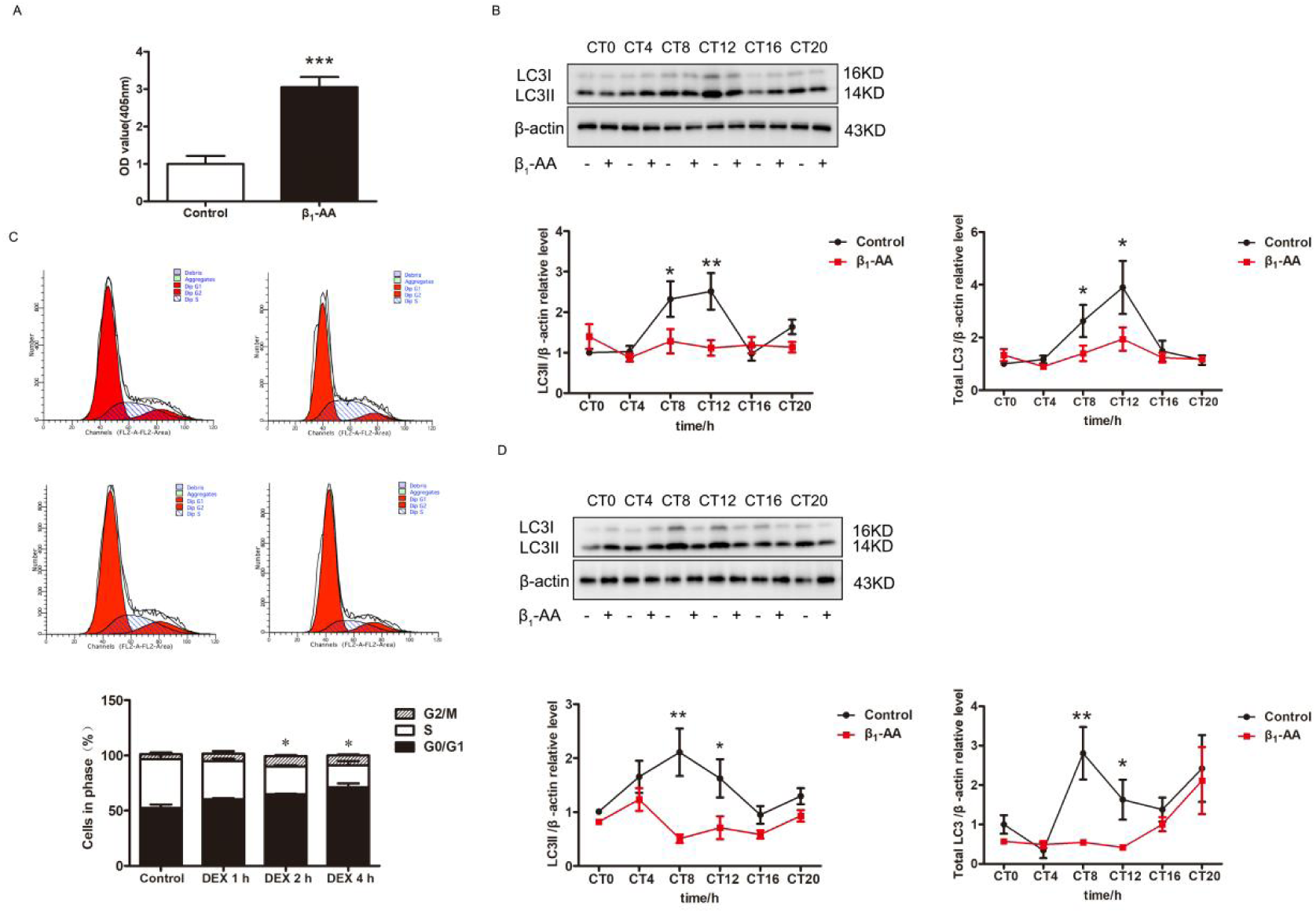
Rhythmic expression disturbance of autophagy marker protein LC3 induced by β_1_-AA A: The serum β_1_-AA antibody content (OD value) of mice in Control group and active immune group was detected by SA-ELISA method; B: Western blot analysis of total LC3 and LC3II expression in myocardial tissue of mice in Control group and β_1_-AA group at different time points (n=14); C: The cell distribution cycle of the Control group was detected by flow cytometry and dexamethasone treatment for 1 h, 2 h and 4 h; D: Western blot analysis of total LC3 and LC3II expression in cardiomyocytes of Control group and β_1_-AA group at different time points (n=7). \**P*<0.05 vs. Control group; \*\**P*<0.01 vs. Control group; \*\*\**P*<0.001 vs. Control group

**Table 1.** Predicting Cyclic Parameters of Myocardial LC3II Protein Expression with JTK_CYCLE Algorithm.

| Sample type | Protein | Group | JTK_CYCLE<br>*P* value | Circadian<br>JTK_CYCLE |
| --- | --- | --- | --- | --- |
| In vivo | LC3II | Control | 0.04 | Yes |
| In vivo | LC3II | $\beta_1$-AA | 0.85 | No |
| In vitro | LC3II | Control | 0.045 | Yes |
| In vitro | LC3II | $\beta_1$-AA | 8.13e-06 | Yes |

Critical value : $P < 0.05$

Then 100 nM Dexamethson was used to induce the synchronization of H9c2 cardiomyocytes. Flow cytometry results showed that the proportion of cells in the DipG1(G0/G1 phase) increased from 52.28% to 71.21% after 4 h treatment with dexamethasone, suggesting that the cells have reached synchronization (Figure 1C). Further detection of LC3II protein expression in cardiomyocytes showed that LC3II protein expression in cardiomyocytes in the Control group had a significant circadian rhythm(JTK_CYCLE ADJ.*P*< 0.05) (Table 1). β_1_-AA could induce the decrease of LC3II protein expression in cardiomyocytes in both CT8 and CT12, with the most significant decrease in CT8. The total LC3 results matched the trend of LC3 II (Figure 1D). Although JTK_CYCLE ADJ.*P* was less than 0.05, LC3II expression decreased at peak value and moved forward in phase under β_1_-AA treatment (Table 1). The above results suggested that β_1_-AA could disrupt the rhythmic expression of myocardial autophagy marker protein LC3.

### 2. β_1_-AA disrupted the rhythmic expression of PER2 protein in cardiomyocytes, mainly by down-regulation of PER2 protein expression

The results of Western blot and JTK_CYCLE analysis showed that the expression of PER2 protein in myocardial tissue of the Control group had obvious circadian fluctuations. PER2 protein expression in myocardial tissue of actively immunized mice was significantly inhibited in CT8, CT12, and CT16 (Figure 2A). Although JTK_CYCLE ADJ.*P* was less than 0.05, indicating a circadian rhythm, the peak value of PER2 expression decreased and the phase moved forward, suggesting that β_1_-AA could lead to rhythmic changes in PER2 protein expression in myocardial tissue (Table 2). The expression of PER2 protein in H9c2 cardiomyocytes in the Control group had obvious circadian fluctuations (JTK_CYCLE ADJ.*P*<0.01) (Table 2). β_1_-AA induced significant inhibition of PER2 protein expression in CT8 and CT16 in cardiomyocytes (Figure 2B). ADJ.*P* value > 0.05 as analyzed by JTK_CYCLE algorithm suggested that PER2 protein expression in cardiomyocytes did not fluctuate in circadian rhythm under β_1_-AA treatment (Table 2). Since both PER2 protein and LC3II protein decreased significantly at CT8 time point under β_1_-AA, the expression of PER2 protein in cardiomyocytes at CT8 time point was further detected by immunofluorescence staining, and the results showed that β_1_-AA caused a significant reduction in the intensity of green fluorescence (FITC labeling) representing the PER2 protein (Figure 2C). These results suggested that β_1_-AA could indeed inhibit the expression of PER2 protein in cardiomyocytes.

**Fig 2.**
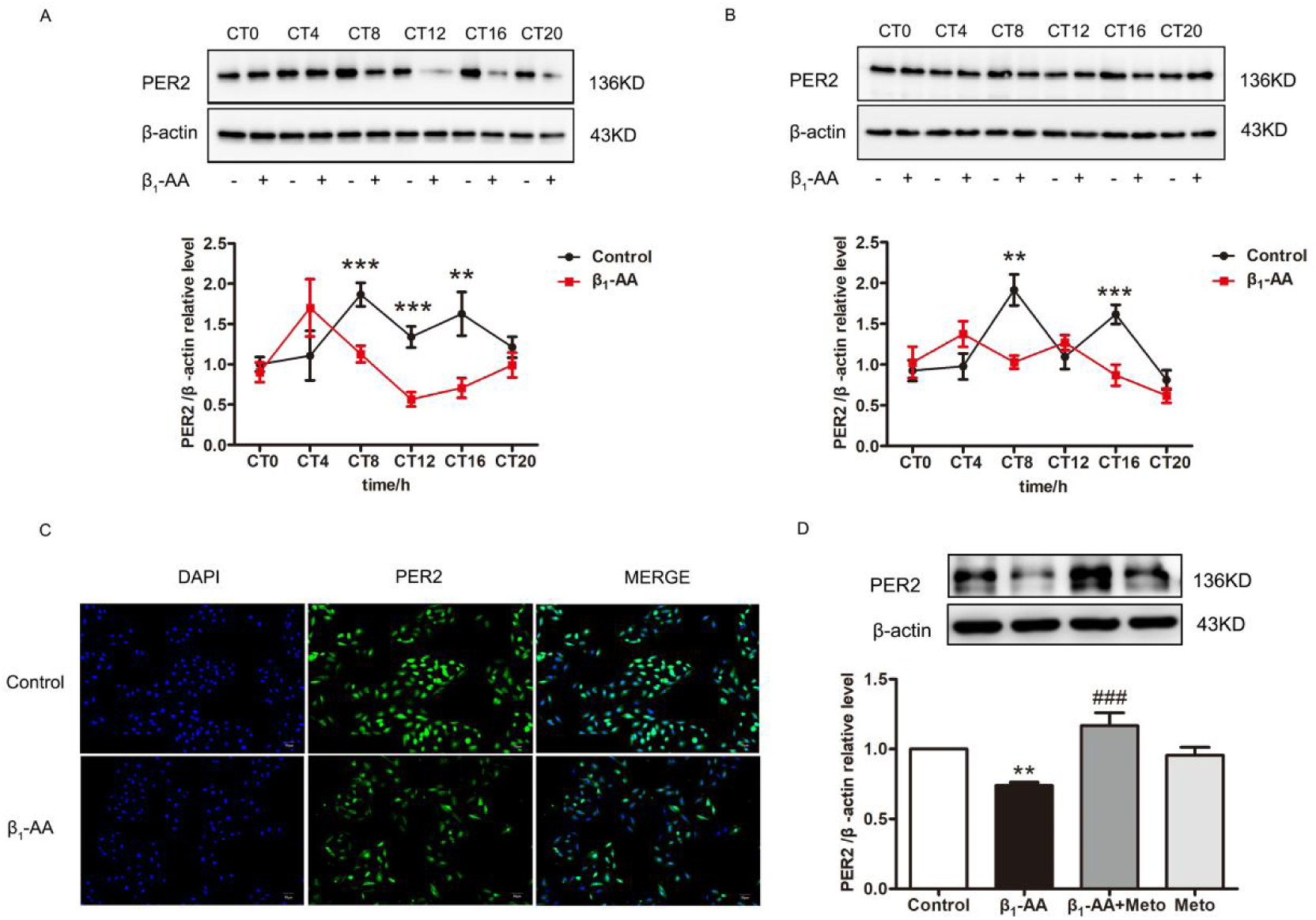
β_1_-AA inhibited the rhythmic expression of PER2 protein A: PER2 expression in myocardial tissue of mice in Control group and β_1_-AA group at different time points (n=6); B: PER2 expression in cardiomyocytes of Control group and β_1_-AA group at different time points (n=12); C: The fluorescence intensity of PER2 protein in cardiomyocytes of Control group and β_1_-AA group was observed (green: FITC-labeled PER2;DAPI: Blue marked nucleus); D: PER2 expression in cardiomyocytes of Control group, β_1_-AA group, β_1_-AA+Metoprolol group and Metoprolol group. \*\**P*<0.01 vs. Control group; \*\*\**P*<0.001 vs. Control group; ^###^*P*<0.001 vs. β_1_-AA group

**Table 2.** Predicting Cyclic Parameters of Myocardial PER2 Protein Expression with JTK_CYCLE Algorithm.

| Sample type | Protein | Group | JTK_CYCLE<br>*P* value | Circadian<br>JTK_CYCLE |
| --- | --- | --- | --- | --- |
| In vivo | PER2 | Control | 0.045 | Yes |
| In vivo | PER2 | $\beta_1$-AA | 0.008 | Yes |
| In vitro | PER2 | Control | 0.014 | Yes |
| In vitro | PER2 | $\beta_1$-AA | 0.262 | No |

Critical value : $P < 0.05$

Further, we selected metoprolol as a specific inhibitor of β_1_-AR to detect the expression level of PER2 protein by setting Control group, β_1_-AA group, β_1_-AA + metoprolol group and metoprolol group. The results indicated that β_1_-AA significantly decreased Per2 protein expression in cardiomyocytes compared to the Control group, while metoprolol reversed this downregulation. Metoprolol alone can also lead to no change in Per2 expression in cardiomyocytes (Figure 2D).

### 3. Decreased PER2 protein expression in cardiomyocytes contributed to the down-regulated LC3 protein induced by β_1_-AA in CT8

To confirm the role of the PER2 protein in the reduction of LC3II expression induced by β_1_-AA, we established cell models for lentiviral Per2 knockdown and overexpression. The optimal multiplicity of infection (MOI) for knocking down Per2 using lentivirus was determined to be 30 (Figure 3A). Stable transgenic cardiomyocytes were obtained after treatment with puromycin for 48 h. Western blot analysis indicated a significant reduction in Per2 expression in cardiomyocytes (Figure 3B). The optimal MOI for the overexpression of Per2 using lentivirus was found to be 20 (Figure 3C). Western blot results showed that Per2 expression was significantly increased (Figure 3D). Subsequently, dexamethasone was treated for 4 h for synchronization, and then β_1_-AA was treated for 8 h. Western blot was used to detect the expression of autophagy marker protein LC3. The results showed that β_1_-AA significantly inhibited LC3II protein expression in cardiomyocytes compared to the Control group. Additionally, LC3II protein expression was further reduced by β_1_-AA following Per2 knockdown. LC3II expression in cardiomyocytes was also significantly reduced with Per2 knockdown alone compared to the Control group (Figure 3E). As shown in Figure 3F, the results showed that overexpression of Per2 significantly reversed this reduction caused by β_1_-AA. Compared with the control group, LC3II expression was significantly increased in cardiomyocytes with Per2 overexpression alone. Similar results were obtained for total LC3 expression (Figure 3G).

**Fig 3.**
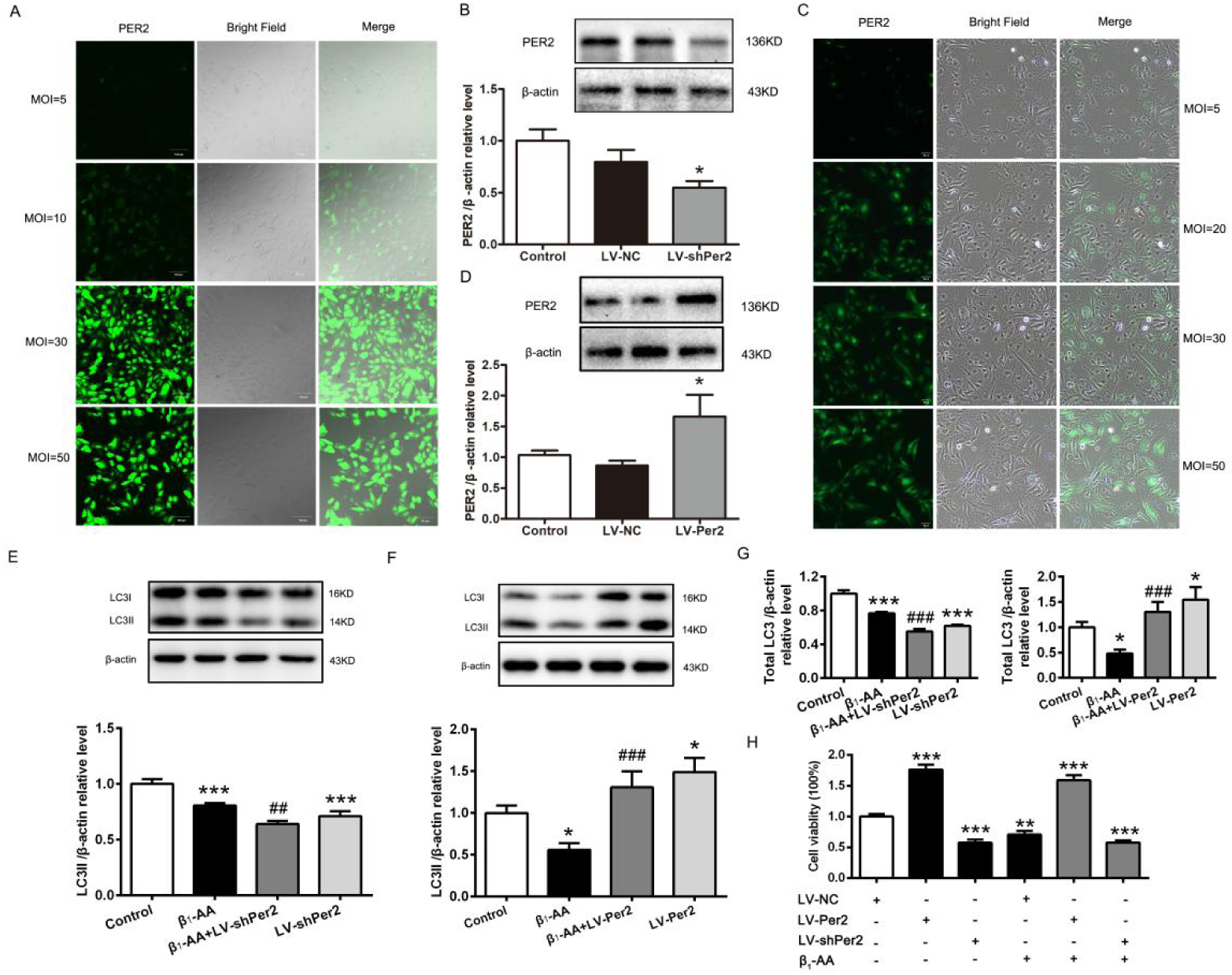
Decreased expression of PER2 protein involved in the downregulation of LC3II protein expression induced by β_1_-AA A: Lentivirus knocked down Per2 expression in cardiomyocytes, set the infection complex number (MOI) gradient and screened the best MOI; B: Western blot to verify whether Per2 was knocked down (n=4); C: Lentivirus infection of cardiomyocytes caused Per2 overexpression, set the infection complex number (MOI) gradient and screen the best MOI; D: Western blot to verify whether Per2 was overexpressed (n=6) ; E: Expression of LC3II in different groups when Per2 was knocked down by Western blot analysis (n=6); F: Expression of LC3II in different groups when Per2 was overexpressed by Western blot analysis (n=5); G: Expression of total LC3 in different groups when Per2 was knocked down (n=6) and overexpressed (n=5) by Western blot analysis; H: The survival of cardiomyocytes after knockdown and overexpression of Per2 was analyzed by CCK8. \**P*<0.05 vs. Control group; \*\**P*<0.01 vs. Control group; \*\*\**P*<0.001 vs. Control group; ^##^*P* < 0.01 vs. β_1_-AA group; ^###^*P* < 0.001 vs. β_1_-AA group

Next, the survival rate of cardiomyocytes was analyzed by knockdown or overexpression of Per2. The results showed that the survival rate of Per2-knockdown cells was significantly decreased, while the survival rate of Per2-overexpression cells was significantly increased. Compared with β_1_-AA alone, the survival rate of cardiomyocytes was still at a low level when Per2-knockdown cells were treated with β_1_-AA. However, overexpression of Per2 could significantly reverse the decreased survival rate of cardiomyocytes induced by β_1_-AA (Figure 3H), suggesting that downregulation of PER2 protein expression in cardiomyocytes induced by β_1_-AA could promote cardiomyocyte death by inhibiting autophagy levels.

### 4. The increased mTORC1 activity in cardiomyocytes was involved in the downregulation of LC3II expression induced by β_1_-AA

In order to explore whether β_1_-AA has an effect on mTORC1 activity, Western Blot analysis was performed to detect mTORC1 activity in cardiomyocytes at CT8. The phosphorylation of mTOR and its downstream molecule S6 is used to represent the activity of mTORC1. The results showed that β_1_-AA induced a significant increase in P-mTOR protein expression and P-S6 protein expression in cardiomyocytes (Figure 4A, 4B). It is suggested that β_1_-AA could promote the increase of mTORC1 activity in cardiomyocytes.

**Fig 4.**
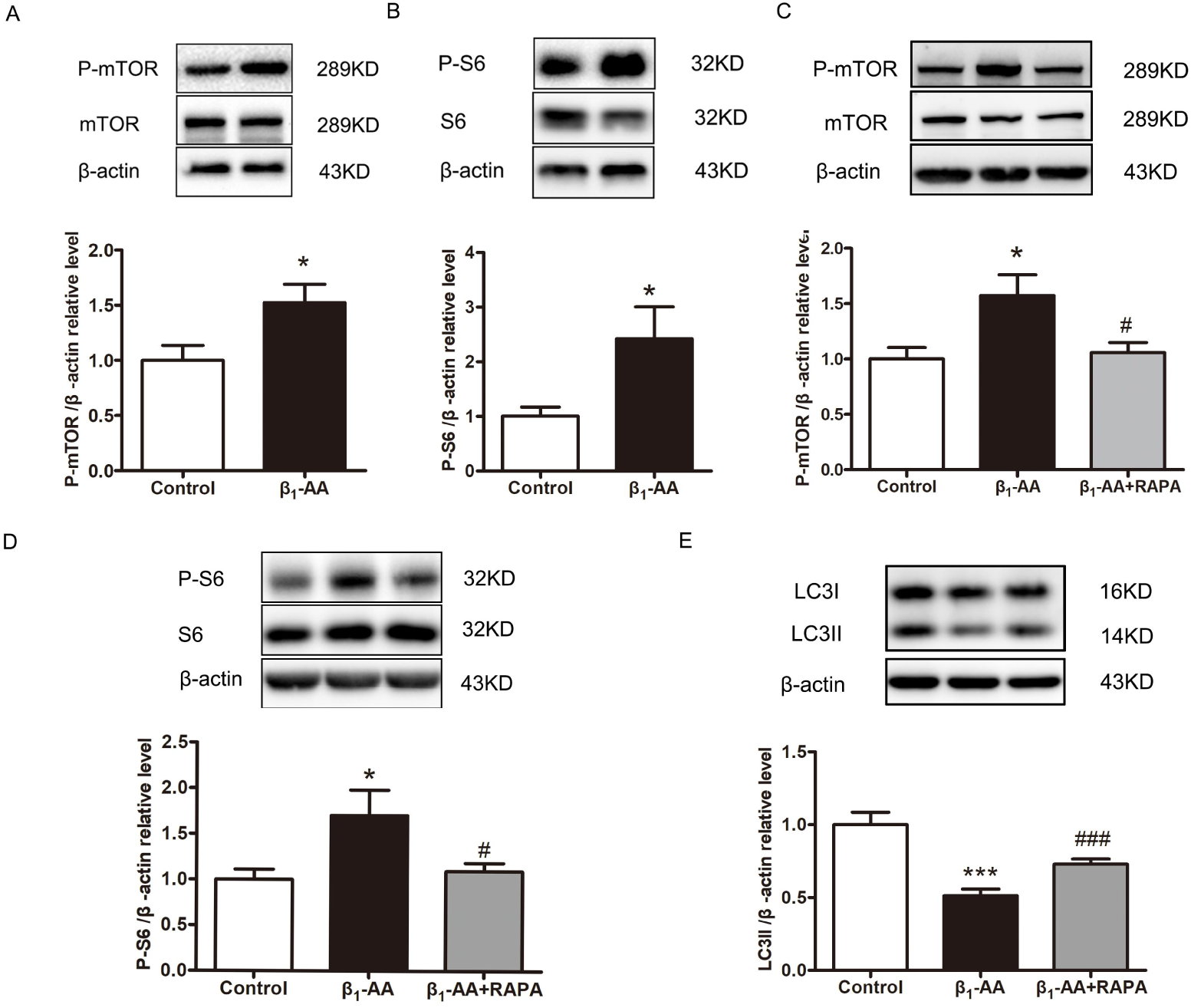
Rapamycin pretreatment improved the β_1_-AA induced decline in the expression of autophagy marker protein LC3II in cardiomyocytes A: Western blot analysis was performed to detect P-mTOR expression at CT8 time point in cells of Control group and β_1_-AA group (n=5); B: Western blot analysis was performed to detect the phosphorylation level of mTORC1 downstream molecule S6 at CT8 time point of cells in Control group and β_1_-AA group (n=4); C: Western blot analysis was performed to detect the phosphorylation level of cardiomyocytes mTOR in Control group, β_1_-AA group and β_1_-AA+RAPA group (n=20); D: Western blot analysis was performed to detect the phosphorylation level of cardiomyocytes S6 in Control group, β_1_-AA group and β_1_-AA+RAPA group (n=9); E: Western blot analysis of LC3II protein expression in cardiomyocytes of Control group, β_1_-AA group and β_1_-AA+RAPA group (n=11) \**P*<0.05 vs.Control group; \*\*\**P*<0.001 vs.Control group; ^#^*P*<0.05 vs.β_1_-AA group; ^###^*P*<0.001 vs.β_1_-AA group

To confirm that mTORC1 activation is involved in the process of inhibition of autophagy levels induced by β_1_-AA. During 4 h of cardiomyocyte synchronization, the mTORC1-specific inhibitor Rapamycin (RAPA) was added to pretreat for 2 h, and then β_1_-AA treatment was performed after the synchronization. The results showed that rapamycin pretreatment significantly inhibited the expression of P-mTOR and P-S6 protein compared with β_1_-AA treatment group alone, suggesting that rapamycin could inhibit mTORC1 activity (Figure 4C, 4D). It was also found that pretreatment with rapamycin followed by β_1_-AA intervention could significantly reverse the β_1_-AA-induced decline in LC3II protein expression in cardiomyocytes (Figure 4E). These results suggested that mTORC1 activation is indeed involved in the inhibition of myocardial autophagy induced by β_1_-AA.

### 5. β_1_-AA promoted mTORC1 activity by inhibiting PER2 expression in cardiomyocytes

In order to investigate whether the decrease of PER2 protein in cardiomyocytes affected the activity of mTORC1 under β_1_-AA, we used lentvirus knockdown or overexpression of Per2 in cardiomyocytes respectively, and detected the phosphorylation level of mTORC1 downstream molecule S6 by Western blot. The results showed that knockdown of Per2 in cardiomyocytes further increased β_1_-AA induced P-S6 protein expression compared with β_1_-AA group and Per2 knockdown alone increased P-S6 protein expression significantly in cardiomyocytes compared with the control group (Figure 5A). Overexpression of Per2 significantly decreased P-S6 protein expression in cardiomyocytes compared with the control group and Per2 overexpression alone decreased P-S6 protein expression significantly compared with the control group (Figure 5B). It was suggested that β_1_-AA promoted the increase of mTORC1 activity by inhibiting PER2 expression in cardiomyocytes.

**Fig 5.**
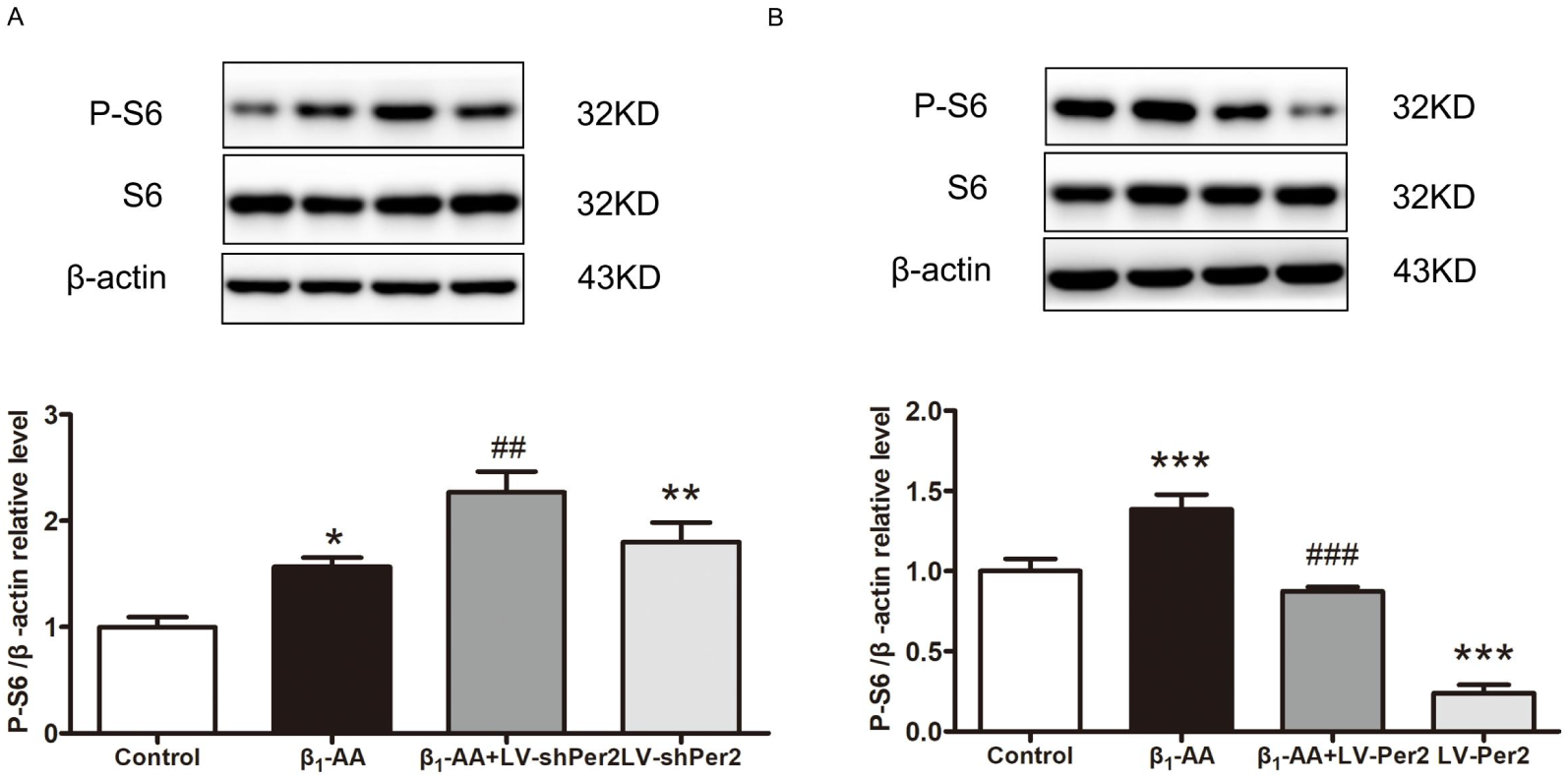
Down-regulated PER2 expression was involved in increased mTORC1 activity in cardiomyocytes induced by β_1_-AA A: The expression level of P-S6 in cardiomyocytes of LV-NC group, LV-NC+β_1_-AA group, LV-shPer2+β_1_-AA group and LV-shPer2 group was detected by Western blot analysis (n=6); B: Western blot analysis of cardiomyocyte S6 phosphorylation levels in LV-NC group, LV-NC+β_1_-AA group, LV-Per2+β_1_-AA group and LV-Per2 group (n=6) \**P*<0.05 vs. Control group; \*\**P*<0.01 vs. Control group; \*\*\**P*<0.001 vs. Control group; ^##^*P*<0.01 vs. β_1_-AA group; ^###^*P*<0.001 vs. β_1_-AA group

**Fig 6.**
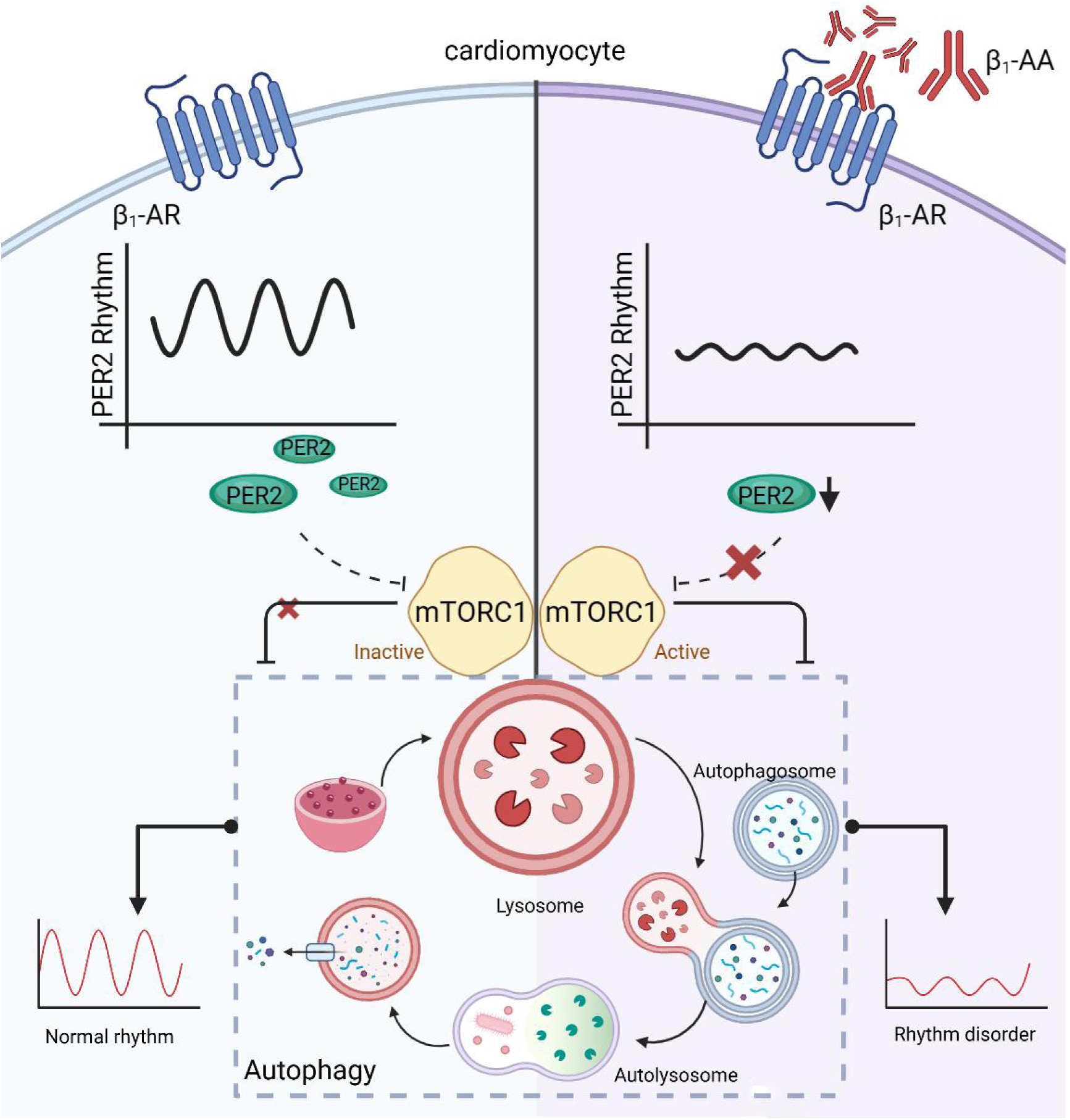
Working model by Created in BioRender.com. is showing that β_1_-AA increase the expression of PER2 in cardiomyocytes. Then, the downregulation of PER2 under the action of β_1_-AA can induce disruption of autophagy rhythm by promoting mTORC1 activation.

## Discussion

Studies have shown that autophagy could maintain cell homeostasis and reduce cell death, and the number of autophagy vacuoles in cardiomyocytes has been found to fluctuate in a rhythmic manner ^[15, 16]^. New statistics showed that most major heart disease outbreaks, such as heart failure, sudden cardiac death, and myocardial infarction ^[17]^, occur in the early morning ^[18, 19]^, when autophagy activity is at a low level throughout the day, suggesting that the occurrence of heart failure may be related to the fluctuation of autophagy rhythm during the day. Studies have found that the disturbance of autophagy rhythm could cause the death of cardiomyocytes, which is one of the important reasons for inducing heart failure ^[9]^. Therefore, it is necessary to find the factors that cause the disturbance of autophagy rhythm of cardiomyocytes in the course of heart failure.

Clinical data showed that an autoantibody, known as β_1_-AA, that persistently activates the β_1_-adrenergic receptor (β_1_-AR) is present in the serum of 40%-60% of heart failure patients. β_1_-AA could continuously activate β_1_-AR on the surface of cardiomyocytes and induce autophagy inhibition, resulting in cardiac cell death and cardiac insufficiency ^[20, 21]^. In this study, we confirmed that β_1_-AA could cause the disturbance of autophagy rhythm of cardiomyocytes. First, we established a β_1_-AA positive mouse model by actively immunized with β_1_-AR-ECII antigen peptide. During this period, mice were subjected to synchronous treatment of LD 2 w to DD 2 w. The specific operation was that the mice were initially kept in the 12-hour light / 12-hour darkness (LD) cycle for 14 days to establish diurnal entrainment, and then kept in constant darkness (DD) to eliminate light interference and observe the endogenous rhythm changes ^[22, 23]^. The results showed that the autophagy rhythm was significantly disrupted in the myocardial tissue of the mice, mainly as the expression of autophagy marker protein LC3II decreased significantly in CT8 and CT12. The JTK_CYCLE algorithm was used to analyze the circadian rhythm parameters of protein expression, and the results showed that LC3II rhythm expression was lost in myocardial tissue of actively immunized mice. Subsequently, H9c2 cardiomyocytes were treated with simultaneous treatment, and dexamethasone (final concentration 100 nM) was added to the petri dish for 1 h, 2 h, and 4 h. Dexamethasone, a synthetic glucocorticoid, activates the glucocorticoid response element on the clock gene promoter. This regulation influences peripheral rhythms, causing most cells to be arrested in the G0/G1 phase, resulting in cell synchronization ^[24,25]^. Flow cytometry was used to detect the proportion of cells in different cell cycles. The results showed that the proportion of cardiomyocytes in G0/G1 phase reached 71.21% after dexamethasone was treated for 4 h, reflecting good synchronization effect ^[26–29]^. Further, autophagy was detected in the synchronized cells, and studies have shown that changes in LC3II levels and total LC3 can be used as indicators of autophagy ^[30]^. After 4 h of dexamethasone treatment, the results found that β_1_-AA could cause the decrease of LC3II protein expression and total LC3 protein expression significantly in CT8 and CT12. JTK_CYCLE analysis showed that although the LC3II rhythm expression was still existed in cardiomyocytes treated with β_1_-AA, the peak value decreased and the phase advanced. These results suggested that β_1_-AA could indeed induce the disturbance of autophagy rhythm of cardiomyocytes. However, how β_1_-AA induces dysrhythmia in autophagy remains unclear.

The biological clock and autophagy are known to be involved in regulating many physiological processes. More and more evidence shows that the circadian clock can regulate autophagy rhythm ^[10,31]^. Among them, the core protein of the biological clock PER2 plays a crucial role in regulating the body’s circadian rhythm. It has been found that isoproterenol, a β-adrenergic receptor agonist, can regulate the rhythmic expression of Per2 ^[32]^. Moreover, overexpression of Per2 has been found to improve myocardial infarction by activating autophagy levels ^[33]^. So, is β_1_-AA induced autophagy rhythm disturbance related to Per2 regulation? The results of this study showed that PER2 rhythmic expression in myocardial tissue of active immune mice was disturbed, which showed the peak value decreased and the phase advanced. β_1_-AA intervention in synchroized cardiomyocytes showed that β_1_-AA could significantly inhibit the rhythmic expression of PER2 protein in cardiomyocytes, especially in CT8 and CT16. JTK_CYCLE analysis showed that the expression of PER2 protein did not have circadian fluctuation. Since both PER2 protein and LC3II protein decreased significantly at CT8 time point under the action of β_1_-AA, CT8 time point was adopted for subsequent experiments. Then, Per2-overexpression and Per2-knockdown lentivirus were used to infect cardiomyocytes and the results showed that in CT8, knockdown of Per2 induced the decrease of LC3 protein expression and cell survival rate was further decreased, while overexpression of Per2 significantly reversed the decrease of LC3 expression and cardiomyocyte survival rate induced by β_1_-AA. These results indicated that β_1_-AA could induce the disturbance of autophagy rhythm by inhibiting the expression of Per2, and eventually lead to the death of cardiomyocytes. Next, we further investigate how Per2 down-regulation is involved in β_1_-AA-induced inhibition of autophagy.

Studies have shown that PER2 protein in mouse liver can regulate autophagy by inhibiting mTORC1 activity ^[34]^. mTORC1 signaling pathway is one of the classical pathways regulating autophagy ^[32]^. The mTOR complex is divided into mTOR complex 1 (mTORC1) and mTOR complex 2 (mTORC2). mTORC1 is composed of mTOR, Raptor, Deptor, mLST8 and PRAS40 subunits. It can inhibit the phosphorylation of UNC-51-like autophagy activating kinase 1 (ULK1) by AMPK, resulting in the failure of LC3II, which is necessary for the formation of autophagosomes, and thus inhibit autophagy levels ^[35]^. More and more studies have shown that abnormal mTORC1 activation is a common phenomenon in many diseases, including heart failure ^[36, 37]^. Thus, we speculate that mTORC1 activation may be involved in the disturbance of myocardial autophagy rhythm induced by β_1_-AA. Western blot analysis showed that β_1_-AA could induce up-regulation of P-mTOR and P-S6 expression levels in myocardial tissue and cells at CT8, suggesting that β_1_-AA could activate mTORC1 activity in cardiomyocytes. In order to further clarify the role of mTORC1 activation in β_1_-AA inhibition of autophagy at CT8, we used mTORC1 inhibitor rapamycin to pretreat cardiomyocytes, and then added β_1_-AA intervention. The results showed that rapamycin could significantly reverse the decreased expression of LC3II protein induced by β_1_-AA. It was confirmed that β_1_-AA could indeed induce autophagy inhibition by activating mTORC1 at CT8. Further, Per2 down-regulation upregulated β_1_-AA-induced P-S6 expression, while Per2 overexpression reversed β_1_-AA-induced P-S6 increase.These results suggested that the down-regulation of PER2 expression induced by β_1_-AA could inhibit autophagy by activating mTORC1 in cardiomyocytes.

In summary, mTORC1 activation caused by down-regulation of PER2 protein expression in cardiomyocytes is an important mechanism of autophagy inhibition induced by β_1_-AA and thus the disturbance of autophagy rhythm. Up-regulation of PER2 protein expression may be an important means to ameliorate the disturbance of autophagy rhythm induced by β_1_-AA and thus improve cardiac function. This study aims to provide experimental basis for the precision treatment of cardiovascular diseases from the perspective of biological rhythm.

## Conclusion

The inhibitory effect of reduced PER2 protein expression on β_1_-AA-induced autophagy rhythm disorder is attributed to increased mTORC1 activity.

## Authorship Confirmation

Peng-Jia Li performed most of the experiments with the assistance of Jia-Yan Feng, Jiao Guo, Jin Xue, Yang Li, and Shi-Yuan Wen conceived the project. Li Wang, Xiao-Hui Wang, Hui-Rong Liu and Peng-Jia Li designed the experiments. Li Wang and Peng-Jia Li analyzed and interpreted the data. Peng-Jia Li and Li Wang wrote the manuscript and revised it critically for important intellectual content. All authors read and approved the final version of the manuscript and its submission.

## Institutional Review Board Statement

All animals from Laboratory Animal Center of Shanxi Medical University [Animal qualification Certificate No. : SCXK (Jin) 2015-0001]. H9c2 rat cardiomyocytes were purchased from Shanghai Cell Bank, Chinese Academy of Sciences.

## Data availability statement

All data generated or analysed during this study are included in this published article.

## Ackonwledgements

The authors would like to give our sincere appreciation to the reviewers for their helpful comments on this article.

